# A mesoscopic simulator to uncover heterogeneity and evolutionary dynamics in tumors

**DOI:** 10.1101/2020.08.18.255422

**Authors:** Jiménez-Sánchez Juan, Martínez-Rubio Álvaro, Popov Anton, Pérez-Beteta Julián, Azimzade Youness, Molina-García David, Belmonte-Beitia Juan, F Calvo Gabriel, Pérez-García Víctor M

**Author notes:** These authors contributed equally to this work (Jiménez-Sánchez Juan, Martínez-Rubio Álvaro).

## Abstract

Increasingly complex in-silico modeling approaches offer a way to simultaneously access cancerous processes at different spatio-temporal scales. High-level models, such as those based on partial differential equations, are computationally affordable and allow large tumor sizes and long temporal windows to be studied, but miss the discrete nature of many key underlying cellular processes. Individual-based approaches provide a much more detailed description of tumors, but have difficulties when trying to handle full-sized real cancers. Thus, there exists a trade-off between the integration of macroscopic and microscopic information, now widely available, and the ability to attain clinical tumor sizes. In this paper we put forward a stochastic mesoscopic simulation framework that incorporates key cellular processes during tumor progression while keeping computational costs to a minimum. Our framework captures a physical scale that allows both the incorporation of microscopic information, tracking the spatio-temporal emergence of tumor heterogeneity and the underlying evolutionary dynamics, and the reconstruction of clinically sized tumors from high-resolution medical imaging data, with the additional benefit of low computational cost. We illustrate the functionality of our modeling approach for the case of glioblastoma, a paradigm of tumor heterogeneity that remains extremely challenging in the clinical setting.

**Author summary:** Computer simulation based on mathematical models provides a way to improve the understanding of complex processes in oncology. In this paper we develop a stochastic mesoscopic simulation approach that incorporates key cellular processes while keeping computational costs to a minimum. Our methodology captures the development of tumor heterogeneity and the underlying evolutionary dynamics. The physical scale considered allows microscopic information to be included, tracking the spatio-temporal evolution of tumor heterogeneity and reconstructing clinically sized tumors from high-resolution medical imaging data, with a low computational cost. We illustrate the functionality of the modeling approach for the case of glioblastoma, an epitome of heterogeneity in tumors.

## Introduction

Discrete mathematical models in cancer track and update individual cells according to a set of biological rules as they interact with other cells and the microenvironment. There is a wide variety of models of this type that include both on-lattice (such as cellular automata) and off-lattice (agent-based) models. With the advent of single-cell resolution technology, next-generation sequencing techniques and the increasing availability of patient data, many mathematical modeling efforts in oncology have been directed towards the use of discrete and individual-based methodologies (see e.g. [1–4] for some reviews). These types of models are being used to address a broad variety of cancer-related problems and some of them are even available as open platforms for broad purpose simulation in cancer [5, 6].

However, discrete individual-based models also have some limitations. They typically incorporate many parameters that have to be obtained from a limited amount of biological information/data. Also, they are difficult to connect with imaging data, since medical imaging has a limited spatial resolution of about 1 mm^3^. Although imaging techniques provide rough information on tumor cell density, metabolic activity, vascular status, and other relevant variables, they do include the details of cellular dynamics within each voxel. Thus, there is room for discrete cell-based modeling approaches beyond classical continuous ones based on partial differential equations but working at the spatial scales at which tumor evolution can be monitored.

From the computational point of view, individual-based models are computationally intensive and suited to describing microscopic scenarios, in-vitro experiments, or even small sections of tissues or model animals. Addressing human tumors in clinical stages normally involves reducing the number of interactions, individuals or processes considered at the microscopic levels, or representing them in a simpler manner. While awaiting progress in computational power that allows for the inclusion of both detailed single cell information and macroscopic simulated tumors, the choice of scale seems to be the dominant factor [7]. As has recently been pointed out [8], few models are able to take into account three-dimensional space, a broad mutational spectrum, mixing populations and reaching clinical or realistic sizes in feasible computational time.

In this paper we put forward a three-dimensional, mesoscale, discrete, on-lattice, multi-compartmental, stochastic approach intended to simulate biological phenomena in clinically-sized tumors. The main element or agent is the cell subpopulation, whose definition is parallel to how species are normally defined in ecological models [9]. Space is discretized in compartments of adjustable size, which allows for comparison with medical imaging data. A compartment is occupied by a number of cell subpopulations with different features, each undergoing dynamics in its spatial location: Growth, interaction with others and spatial spreading. This intermediate scale allows for the integration of detailed biological data and for computationally feasible simulations up to the macroscopic, whole-tumor scale.

The global evolution of the tumor is driven by the dynamics at each lattice position which, in turn, is governed by the behavior of the different cell subpopulations, which are subject to four fundamental biological processes: reproduction, death, mutation and migration. These four processes, stemming from the basic hallmarks of cancer [10], occur stochastically in a synchronous manner, meaning that at each time step the population is updated according to the probabilities computed in the previous step. As a result, the system moves from a very simple initial state to a fully grown, realistically sized, heterogeneous tumor, reconstructing its entire natural history. Benefiting from an optimized computational time, this model can constitute a platform for hypothesis testing or therapy simulation or any issue that the medical community, ideally involved in this process, would consider of interest.

As a bench test to assess our model’s performance and versatility in tumor modeling, we also show a detailed application of the mesoscopic model to glioblastoma (GBM), one of the most aggressive tumors, which also epitomizes intratumor heterogeneity and enhanced phenotypic adaptation capacity [11], with no major improvements in outcome since the establishment of the standard Stupp protocol [12]. With the notable example of the TCGA genomic characterization of GBM [13], information about the mutational spectrum of this tumor [14] and the relative frequencies [15], coexistence [16] and exclusivity [17] is now available, sometimes including spatial information [18] and reconstructions of its evolutionary history [19, 20]. Imaging data is also increasingly available, providing valuable details related to geometry, shape, size, regularity, and it is also used in biomarker identification [21–23]. It is therefore an ideal scenario for testing and calibrating a model for tumor growth and diversification that includes molecular information and reaches clinical sizes.

## Materials and methods

### Computational model

#### 3D Lattice

Space is discretized as a hexahedral mesh consisting of *L*_*x*_ × *L*_*y*_ × *L*_*z*_ spatial units (voxels) or compartments, with *L*_*i*_ being the number of compartments per spatial dimension. Both the volume and number of compartments in the grid are adjustable parameters. Since high-resolution imaging (e.g. 3D magnetic resonance imaging T1/T2/FLAIR sequences) voxel size is around 1 mm^3^, we chose that voxel size for the specific examples. No-flux boundary conditions were set. However, we chose lattices large enough that simulations typically do not reach the boundaries of the domain.

#### Clonal subpopulations

Each voxel contains cells that undergo different cellular processes: division, migration, death and mutation. A cell belongs to a clonal subpopulation that is defined by a set of traits. Traits are represented by a vector 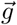 of length *G*, 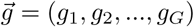, where *G* is the number of traits or alterations and *g*_*i*_ can take the values 0 or 1. This value can be interpreted as two possible expressions for a trait, or presence or absence of a mutation, as typically done when treating species in ecological models [9]. The clonal subpopulation then constitutes the basic unit or agent in the computational model, trading cell resolution for feasible simulation time and achieving clinical tumor sizes. The set of traits that represents a subpopulation determines the rate at which cells undergo biological processes, so that cells from a given subpopulation on a given voxel will behave in the same way, except for stochastic noise intrinsic to cell processes (biological instances of this are differential gene expression or variable mitochondrial content). Populations with more advantageous traits will be more likely to become fixed in the tumor, especially once they achieve a large cell number, whereas at early stages (low cell numbers) genetic drift will be more important. The processes are modeled in such a way that cells grow exponentially when there is enough empty space and slow their growth as the voxel becomes crowded. Migration is also influenced by voxel occupation as described below.

#### Cell division

Each clonal subpopulation has a different reproduction probability depending on its traits and its local (voxelwise) environment. Mathematically, this probability is expressed as:

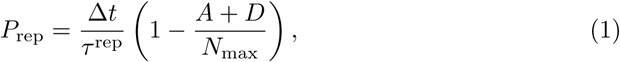

where Δ*t* is the time step considered, *A* and *D* are the total active and necrotic population in each voxel respectively and *N*_max_ is the local carrying capacity. The probability is modulated by the relationship between time step and process characteristic time; the probability of a process to occur increases with the time step. The reproduction probability decreases with occupation, simulating competition for space and resources. *τ*^rep^ is the part of the probability that depends on each subpopulation’s traits. It has units of time and is computed as

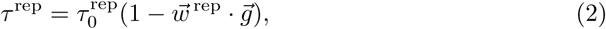

where the first term, 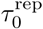, is a basal reproduction time, assigned to the wild type, and the second term represents how this basal rate is modified by the different alterations that the cell can undergo. Also, 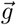 is the trait vector and 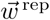 is a vector having the length *G*, with *w*_*i*_ representing the degree to which an alteration modifies the respective rate. It satisfies |*w*_*i*_| < 1 and Σ_*i*_ *w*_*i*_ ≤ 1.

Taking into account the whole clonal subpopulation, we replace the time-consuming Montecarlo process with a direct sampling from a binomial distribution *X* ~ B(*A, P*_rep_) with *A* being the number of active cells of the subpopulation in the previous iteration.

#### Cell migration

The migration process occurs in two differentiated steps. Firstly, the number of cells leaving the current voxel is computed. They are then distributed into neighboring voxels, taking into account their relative distances. The probability of migration for a single cell is given by

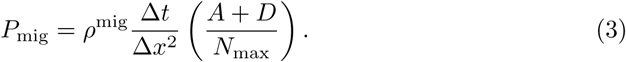

Eq (3) has a similar structure to the reproduction process in Eq (1) albeit the trait-dependent term *ρ*^mig^ has the units of a diffusion coefficient instead of a being the inverse of a characteristic time, and differs from the impact of each alteration on the process, 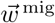. We include here the spatial step Δ_*x*_, by analogy with the discrete Laplacian operator in space, in order to keep the probability adimensional. We calculate *ρ*^mig^ in the following recursive form, similar to (2), as

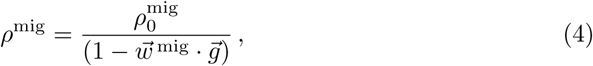

where 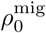 is a basal diffusion constant corresponding to the wild type modulated by subsequent alterations. Notice that the product between 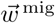 and 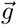 is moved to the denominator in order to keep the same structure as in the other basic processes. In contrast with the division probability, the migration probability increases with the occupation of the voxel, since a cell is more likely to migrate if there is more competition for space and/or resources. Necrotic cells do not migrate, but they do occupy space, and thus must be taken into account. The total number of migrating cells is again sampled from a binomial distribution *X* ~ B(*A, P*_mig_).

As for the destination of each cell migrating from a voxel (*x, y, z*), we considered a Moore neighborhood 𝓜_*x,y,z*_ in three spatial dimensions. In this way, each migrating cell has 26 possible desvtinations. Each destination has an associated probability with weighting factors 1, 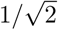 and 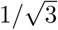 when voxels share a face, edge or vertex with the central voxel respectively. The destination is then computed by sampling migrating cells from a multinomial distribution with the 26 probabilities. Performing the migration in this way reproduces a diffusive process, in which migration depends on cell density gradients and distances.

#### Cell mutation

Mutation is the mechanism for the diversification of the population, and is considered here in a broad sense as a change in one of the characteristics of the cell, be it genetic or phenotypical. The mathematical formula used to compute the mutation probability is

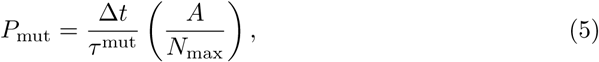

so that mutations depend only on *A*, the number of cells in a given subpopulation; *N*_max_, the carrying capacity, and *τ*^mut^, the characteristic time that includes trait effects as explained above. This probability measures the chance of a mutational event occurring, not the specific alteration; this is later randomly selected from the subset of *G*, representing non-altered traits. Since the time scale of mutational events is much longer than that of cell division or death, mutations are assumed to occur in individual cells (Bernoulli process) instead of sampling them from a binomial distribution.

#### Cell death

The form of the cell-death term is similar to the proliferation term, with the difference that the probability of death increases with occupation. The expression for the probability of death is then given by

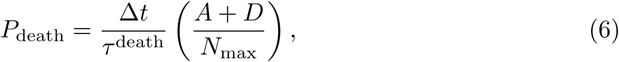

where *τ*^death^ is a typical cell lifetime. The reason for this dependence is that in aggressive tumor cells induce a damage to the microenvironment that leads to their own death. In the example to be studied in this paper, glioblastoma, it is well known that this happens through the secretion of prothrombotic factors that induce microvessel failure and the formation of necrotic areas [24].

#### System updating

Let 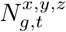 be the number of cells in subpopulation *g* at voxel (*x, y, z*) and time *t*, for *g* = 1, …, 2*^G^* and *x* = 1, …, *L*_*x*_, *y* = 1, …, *L*_*y*_, *z* = 1, …, *L*_*z*_. The number of cells at the discrete time *t* + 1 is computed using the equation

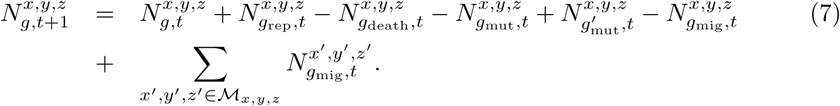

Eq (7) governs the updating of the cell number of each clonal population. It includes the positive contributions of newborn cells 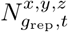, mutations from other clonal populations that come to the one evaluated 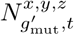 and cells in the same subpopulation migrating from other voxels in the Moore neighborhood 𝓜_*x,y,z*_ of current point 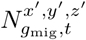. Cell numbers decrease by subtracting dead cells 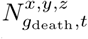, cells mutating to different clonal subpopulations 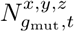 and cells migrating to surrounding voxels 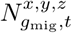. Contributions are randomly sampled at each time step from the respective distributions as explained above, taking into account the probabilities calculated for each process. The three-dimensional structure of our model resembles that of a multicompartmental cellular automaton, with many individuals per grid site. In that context, it is well known that the way the grid is updated may have an impact on the behavior and end state of the system [25]. We opted here for uniform time discretization and synchronous updating, computing the change in population in each voxel at each time step according to the stochastic biological rules. Then we made use of an auxiliary structure, to transfer all the changes to the next time step, thus avoiding updating artifacts. This process was repeated for the whole time of the simulation. A summary of the algorithm can be found in Fig 1.

**Fig 1.**
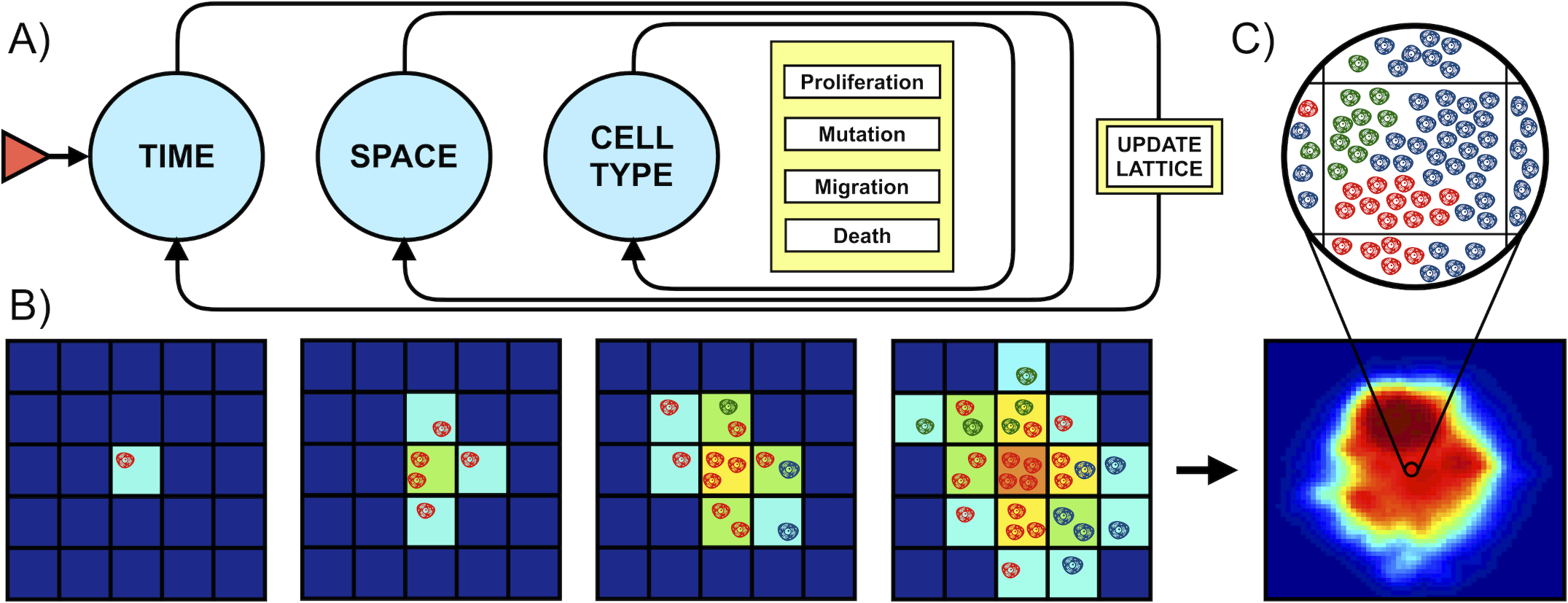
Algorithm. **A)** Basic algorithm (for implementation purposes). Initialization requires creating a 3D grid, specifying the final simulation time, defining subpopulation traits and setting the initial state. Temporal iterations are then carried out until the end time is reached. At each time step and each voxel, all clonal populations are updated. This updating involves calculating how many cells will proliferate, migrate, mutate or die. When all populations at all voxels have been evaluated, they are updated synchronously. **B)** Two-dimensional example of model behavior. Synchronous updating results in population increasing, diversifying and spreading at each time step, with probabilities computed according to the biological rules. Compartment color indicates occupancy. Cell color indicates cell type. **C)** Bottom image is a slice of an actual simulation, with colors indicating occupation. Each voxel contains a variable number of cells and subtypes as depicted above.

### Estimation of parameters

Since one of our goals is to apply the mesoscopic simulator to the case of GBM as a benchmark test, we used the sizes typically found in the clinical setting for the tumor maximum sizes, which are around 100 cm^3^. Hence, we selected *L* = 80 to make these sizes attainable. The time step was fixed at 4 hours and simulation ended when the tumor reached 100 cm^3^. From typical cell sizes [26] we estimated the carrying capacity of a single voxel *N*_max_ to be 2 × 105 cells.

Sequencing studies of GBMs [13, 15] reveal that most of the mutations found in this disease cluster around three main pathways: RTK/PI3K/RAS, RB and p53. Each of these pathways has different key alterations and frequencies. We therefore considered a set of three possible alterations characterizing populations (*G* = 3), so that 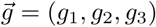. This means that there are eight possible cell subtypes (2^*G*^), depending on all possible combinations of altered pathways (Fig 2).

**Fig 2.**
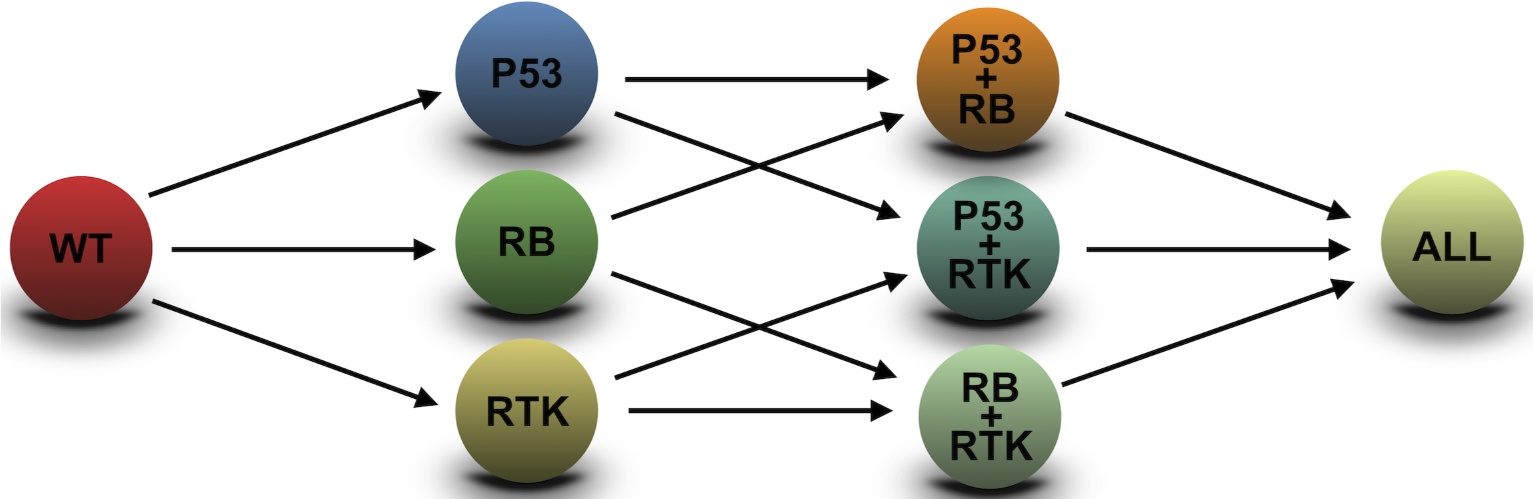
Mutation tree used in this paper. Relationships between the eight possible genotypes, according to the three alterations selected. Each clonal type can emerge from several ancestors by various alterations. Depending on the mutational history, tumors follow different paths on the mutation tree.

The choice of division, death, mutation and migration basal rates used a Bayesian criterion. To obtain an initial coarse estimate of the parameter ranges, the most straightforward way is to use imaging data from real GBMs. Basal reproduction and death rate (Table 1) were estimated from papers using exponential/gompertzian growth laws to fit GBM growth curves [27, 28]. Basal migration parameters were estimated from [28]. Since monitored tumors already carry an unknown mixture of alterations, these numbers should be taken only as rough estimates. Basal mutation rates were estimated from the known values per base and generation for each altered pathway [29–31]. To refine those ranges, thousands of simulations were run with input parameters randomly sampled from previous ranges. Simulations whose tumor lifespans were substantially longer than those typical of real GBMs were rejected. Simulations whose tumor lifespans were close to those of real GBMs were accepted. Basal rate ranges were thus constructed on input parameters from accepted simulations.

**Table 1.** Parameter values for simulations of GBM. Basal rates refer to cells with no mutations. Weights specify how each mutation affects the basal rate.

| Alteration | Reproduction | Migration | Mutation | Death |
| --- | --- | --- | --- | --- |
| RTK | 0.32 | 0.65 | 0.18 | -0.15 |
| RB | 0.28 | 0.05 | 0.18 | -0.05 |
| p53 | 0.25 | 0.05 | 0.32 | -0.45 |
| Basal | 80-250 h | 0.0033-0.0125 $\text{mm}^2/\text{h}$ | 80-240 h | 80-400 h |

The impact of a mutation on the basal rate (weight *w*_*i*_) is less straightforward to determine, as there is no experimental way of estimating how a single mutation affects a given cellular process in living tissue. We followed a similar procedure to the above: for proliferation, for example, by having estimates of maximum and minimum tumor doubling times we can compute the maximum degree to which it is affected by mutations and assign to each alteration a fraction of that modification. This assignment was carried out using available biological information on the functions and processes associated with each pathway. For instance, RTK alterations typically promote proliferation and migration, and p53 promotes avoidance of death, as well as increased genetic instability and thus a higher probability of mutation [32]. We thus ensured that populations undergoing all mutations did not reach unrealistic proliferation rates.

Many sets of weights were tested. Here we show the results using the same set across all simulations so as to make the study reproducible. Further work on this model will involve the estimation of a realistic weight set by more elaborate means. The choice of weights is shown in Table 1 as well as the chosen ranges for the values of the basal rates. Notice that these rates are associated with cellular processes; whole-tumor rates emerge as a result of combined cellular processes. Cellular traits were randomly sampled from the range of allowed basal rates for each simulation. This provided variability between individual simulations and allowed us to assess the robustness of the model’s behavior.

### Macroscopic tumor measures

We will use a set of measures to quantitatively compare tumor longitudinal dynamics with the solutions of the discrete simulation model. We list them here and give precise definitions.

#### Heterogeneity

Diversity indexes are used in ecology to track genotype heterogeneity [33]. Two of the most frequently used are Shannon entropy and the Simpson index. The Shannon entropy quantifies the uncertainty in predicting the species of a selected individual in a population.

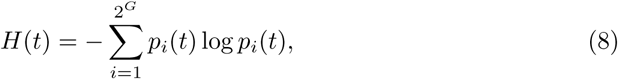

where *p*_*i*_(*t*) is the proportion of individuals from species *i* in the tumor at time *t*:

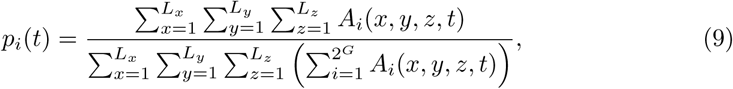

where *A*_*i*_(*x, y, z, t*) is the number of active cells from species *i* in voxel (*x, y, z*) at time *t*. The Simpson index quantifies the probability of picking two individuals at random from the same subpopulation:

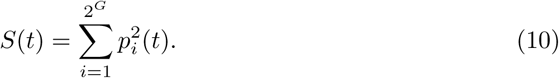

A Shannon entropy equal to 0 means that all cells in the system belong to the same subpopulation, so there is no uncertainty in predicting cell type. A higher Shannon index means higher uncertainty and thus higher heterogeneity. A Simpson index of 1 indicates that all cells belong to the same type, while a value of 0 shows that there are no cells of equal type. In this study we were interested in heterogeneity dynamics of the whole tumor, so we considered Shannon entropy and the Simpson index integrated over all space, as functions of time. Using these indexes we can infer whether several cell populations with different mutational profiles coexist within the tumor, or a single cell population dominates over the others.

#### Volumetric and Morphological measures

Let 𝓥 be the set of voxels that have reached more than 20% of their carrying capacity, considering both living and necrotic cells. Let 𝓥_CE_ be the subset of 𝓥 consisting of voxels in which active cells alone have reached more than 20% of the carrying capacity. Let 𝓥_I_ be the complementary subset 𝓥_I_ = 𝓥\𝓥_CE_. Let us define the number of elements in each set by *N*_*CE*_ = |𝓥_*CE*_|, *N*_*I*_ = |𝓥_*I*_| and *N*_*T*_ = |𝓥|. Note that, because of this definition 𝓥_*CE*_ ⋂ 𝓥_*I*_ = ∅. Then, if individual voxel volume is *V*_vox_, we define the contrast-enhancing (*V*_*CE*_) and inner or necrotic (*V*_*I*_) volumes as

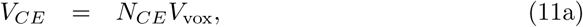

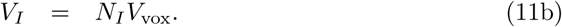

Contrast-enhancing volume is associated with active tumor regions, while inner volume represents the necrotic core. Both quantities can be obtained from computer simulations of our mathematical model. The sum of both magnitudes represents the whole tumor volume *V*. The surface 𝓢 enclosing 𝓥, and its measure *S*, were obtained using the marching cubes method, seen in [23], to resemble the method used to extract this feature from MRIs.

We also defined the mean spherical radius (MSR), as the radius of a sphere having the same volume as the tumor, i.e.

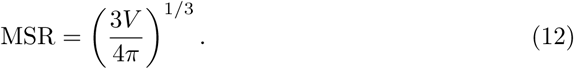

In addition to the volumetric measures we also employed several morphological descriptors that have been found to have prognostic value in different tumor types. They are active tumor spherical rim width (*δ*_*s*_) and surface regularity (*S*_reg_). The first one is obtained from MRIs as an averaged distance between the contrast-enhancing volume and the necrotic core [34]. It can be computed from the volumes through the formula

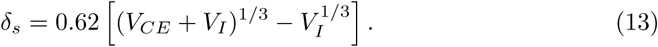

This biomarker has been found to have prognostic value for GBMs using both MRI [22, 34] and PET [35] images.

To quantify the surface regularity we used a dimensionless ratio defined as the relation between the total volume tumor *V* and the volume of a sphere with the same surface *S* [23]:

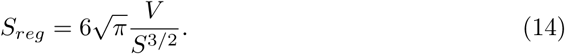

The closer this ratio is to 1, the more similar to a sphere a tumor will be (more regular). When *S*_*reg*_ approaches zero the tumor will be highly irregular, resembling a fractal-like structure. This parameter receives different names in the literature and has been found to have prognostic value in lung cancer [36, 37], head and neck cancer [38, 39], esophageal cancer [40], breast cancer [41], lymphoma [42], and glioma [23, 35].

#### Metabolic measures

Metabolism in tumors can be imaged using positron emission tomography (PET). Different metrics of tumor metabolism are routinely obtained in the clinic. They include the maximum standardized uptake value (SUVmax), the metabolic tumor volume (MTV), and the total lesion activity (TLA). It has been pointed out that TLA and MTV are related through a scaling law providing a surrogate for tumor aggressiveness [43]. The mathematical expression that defines this law is

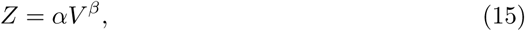

where *Z* is an observable quantity associated with tumor metabolic rate, *V* is a measure of the size of the system (typically mass or volume) and *β* is the scaling exponent. Metabolic scaling laws have implications for how tumors grow [44]. Sublinear exponents *β* < 1 lead to bounded growth, while superlinear exponents *β* > 1 lead to an explosion in size in finite time. Malignant tumors appear to follow the latter trend and the deviation from the reference scaling exponent has prognostic value [43]. In real tumors, the scaling exponent can be computed as

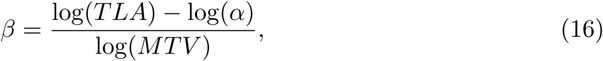

In our simulations, *TLA* can be obtained from the proportion of proliferating cells, since tumors use most of their energy in proliferation, and *MTV* from spatial on-lattice measures.

### Tumor growth law

The search for the mathematical equations that govern tumor growth has been a constant in the history of mathematical oncology [45, 46]. Several attempts have been made to fit tumor growth laws to longitudinal volumetric data, including glioblastoma [27, 47]. Here, we aim to reproduce this for the simulated tumors. We select exponential, gompertzian and radial growth, all of which have been analyzed in the previously cited studies. Motivated by the metabolic analysis described above we also tried to fit to a power law, a variation of the von Bertalanffy equation and another usual candidate for tumor growth law. The equations to be fitted are therefore:

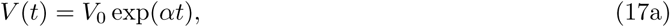

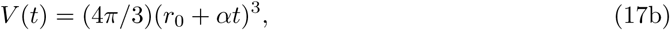

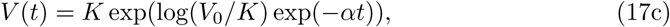

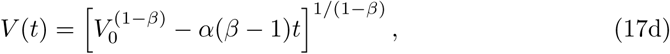

where *V*_0_ is initial volume and *r*_0_ the corresponding mean initial radius, *α* denotes respective growth rates, *K* is the carrying capacity in the Gompertz model and *β* is the scaling exponent in the power law. We used Matlab function lsqcurvefit to obtain fitted parameters and root-mean-square error in order to compare goodness of fit. Initial value was fixed to that of the simulated tumor and the carrying capacity in the Gompertz model was assigned an upper bound of 1400 cm^3^ (average cranial capacity). For the power law, we tried different scaling exponents *β* and selected the one providing the best fit.

### Survival analysis

The measures explained above have prognostic value in real GBMs. We performed a Kaplan-Meier analysis over sets of simulations to evaluate the model’s ability to reproduce this behavior. We set diagnostic time arbitrarily to be the time at which the tumor reaches 1 cm^3^ plus a randomly sampled time from an exponential distribution in order to account for variability of tumor size at diagnosis. Simulations ended when the tumor volume was larger than 100 cm^3^.

Having a survival time for each patient, we performed a search over all possible splitting thresholds. A splitting threshold was used to separate simulations into two groups: tumors with prognostic measurements higher than the splitting threshold will be identified as belonging to an aggressive group, while tumors with prognostic values lower than the splitting threshold will constitute the benign group. The splitting threshold with the lowest p-value was selected, provided that the largest group was not more than 3 times the size of the smallest group.

To obtain the scaling law exponents, we performed a linear regression over log-log data of tumor volume and activity at the time of diagnosis, and obtained the scaling law exponent *β*, which is equivalent to the slope (Eq (16)). Given this information, tumors falling below the regression line were described as hypoactive, while tumors falling above the regression line were called hyperactive. These two groups were used to construct the survival curves. Statistical significance was computed using the log-rank test. Time separation between curves at median survival was also calculated.

Results from the survival analysis may depend on the stochastic sampling of diagnostic times. In order to provide robust results, a Kaplan-Meier analysis was repeated 1000 times with different seeds, thus providing different diagnostic times for each simulation. For each repetition, the p-value was measured, to check whether results were consistent and independent of the random number generated.

### Software

The model was coded both in Matlab (R2018a, The MathWorks, Inc., Natick, MA, USA) and Julia (version 1.1.1). The main workspace and simulation sections were coded in Julia, while data analysis and plotting were coded in Matlab. Simulations were performed on three different machines: a 12-core 32 GB RAM 2.7 GHz Mac Pro, a 6-core 16 GB RAM 3.7 GHz custom-built computer, and a 2-core 8 GB 2 GHz MacBook Pro. Computational cost per simulation was of the order of 3-5 minutes, depending on the machine used and the simulation parameters.

### Tumor rendering

Three-dimensional tumor volumes were rendered for visualization purposes. Total tumor volume was smoothed with function `smooth3` with default settings. Isosurfaces were then computed with the `isosurface` function with isovalue 0.6. Figures were created with the `patch` command and using Matlab’s `Zbuffer` renderer. Additional settings included default `camlight` and `phong` lighting.

## Results

### Simulated tumors recapitulated known GBM timescales and resemble clinical imaging data

To simulate the basics dynamics displayed by the model, we ran a set of 100 simulations starting from one wild-type cell (i.e. without mutations) placed at the center of the spatial domain. Each simulation had a different set of basal rates, sampled randomly from the ranges specified in Table 1 as discussed previously. This allowed us to study variability in the tumor dynamics. Tumors evolved according to the rules explained in the previous section and the volumetric, morphological and metabolic macroscopic variables were tracked as discussed in the ‘Methods’ section. Results are shown in Fig 3. The range of basal rates considered allows for the appearance of tumors with long inception (up to 13 months) as well as rapidly growing cancers (less than 5 months combining inception and growth), both in terms of MSR (Fig 3E) and volume (Fig 3F). These values are in agreement with clinical experience since treated GBM patients have a median survival time of around 15 months [12].

**Fig 3.**
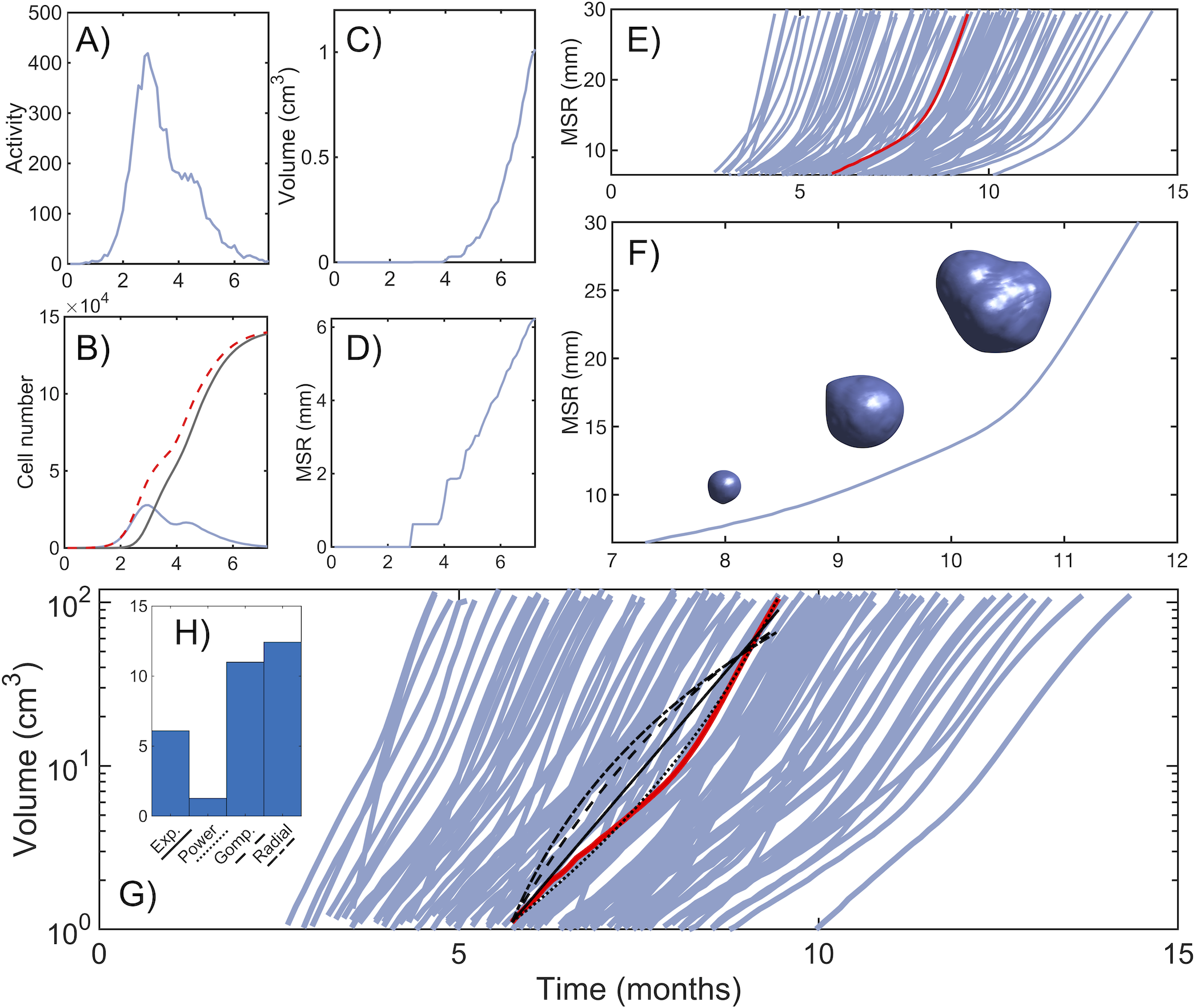
Longitudinal tumor growth dynamics. **A)** Time dynamics of the number of newborn cells at the central voxel (surrogate for tumor activity). **B)** Number of cells at the central voxel: total (dashed red), active (blue) and necrotic (grey). **C,D)** Tumor volume and MSR during the initial stages, starting from a single cell at the center of the lattice until tumor reaches 1 cm^3^. **E)** Dynamics of MSR for 100 simulations. Median run is shown in red. Time span shown starts when tumor reaches 1 cm^3^ in volume (equivalent to 6.2 mm of MSR) and ends after reaching 100 cm^3^. F) Example of tumor dynamics of the MSR and rendered 3D tumor shape for three different times (8.5, 10, 11.5 months). Basal rate parameters for simulation in this figure are *τ*^rep^ = 216.5 h, *τ*^death^ = 112.7 h, *τ*^mut^ = 200.4 h, 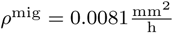. **G)** Dynamics of tumor volume for 100 simulations. Black lines represent different fits of the median run (solid red line): Exponential (solid), power law with *β* = 1.2 (dotted), gompertz (dashed) and radial (dashed-dotted). **H)** Root-mean-square error (RMSE) of each fit.

The filling of a single voxel follows a dynamics resembling logistic growth. An initial fast-growing phase is followed by a peak in the number of active cells, which begin to decline due to saturation-induced death and migration to surrounding voxels (Fig 3B). While the total number of cells (both active and necrotic) tends to the carrying capacity in the long term, active cells decline to zero steadily as they die due to damage to the microenvironment. This can be seen in Fig 3A, where newborn cells reach a peak in activity, early in growth, and then decline steadily. The logistic-like dynamics within individual voxels does not imply that the global growth of the tumor is also bounded. This is because the available physical space around populated voxels allows for sustained growth in tumor volume (Fig 3C). We tried to fit this sustained growth to different growth laws as explained in the ‘Methods’ section (Fig 3G). The best fit for the median run according to the RMSE corresponded to a power law with scaling exponent of *β* = 1.21 (Fig 3H).

Since space is discretized in hexahedral units, and resolution is low for small tumors, volume and surface calculations are not precise until tumor cells have spread to a large group of voxels. This is shown in Fig 3D, and especially affects those measures that approximate the tumor to a sphere (MSR, rim width, surface regularity). Because of this, tumor measures are only reliable when tumor volume exceeds 1 cm^3^. Once this detectable size has been reached, MSR behaves linearly up to a point where there is acceleration in growth, due to the appearance and competition posed by more aggressive genotypes that increase the global growth rate. This is more clearly seen in Fig 3F, which shows the evolution of the MSR along with three-dimensional reconstructions of the tumor for a typical run. The sharp increase in size occurs during the last two months of the disease, in agreement with the known lethal progression of these tumors [48].

Fig 4 shows two-dimensional slices of six simulated tumors, and one of a post-contrast pre-treatment T1-weighted MRI scan of a GBM patient. In this type of image, white areas correspond to regions where an intravenously injected gadolinium contrast is released into the tissue. The reason is that tumor blood vessels are less stable and lack functional pericytes. Thus, this marker is a surrogate of tumor cell density, leading to more abnormal vessels and suggesting that brighter areas would correspond to higher tumor cell loads. However, above a certain density, tumor cells damage the microenvironment and the secretion of prothrombotic factors leads to massive local cell death [24]. Inner dark regions represent necrotic tumor areas calculated as explained in ‘Methods’. The presence of a tumor-enhancing rim enclosing a highly irregular shape, which is a characteristic hallmark of GBM, is also captured in our simulations, and represents a region of highly proliferating cells [49].

**Fig 4.**
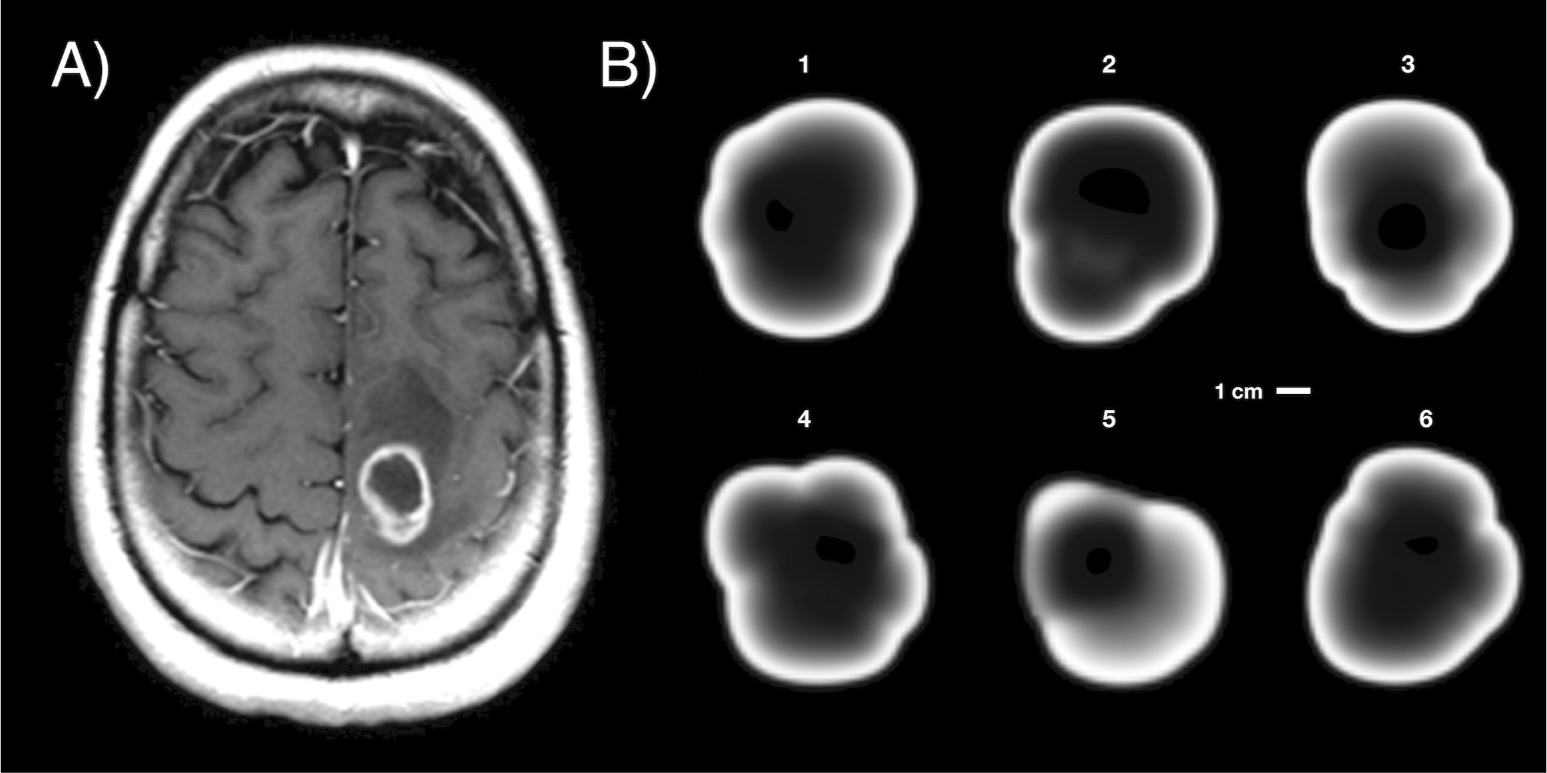
Tumor slice and simulated profiles. **A)** Two-dimensional slice of a T1-weighted post-contrast MRI scan of an actual GBM. **B)** Two-dimensional slices of different simulated tumors. Each simulated tumor corresponds to the final state (100 cm^3^) of a different running of the model. Basal parameters for each simulation are: **1)** *τ*^rep^ = 242.6 h, *τ*^death^ = 213.27 h, *τ*^mut^ = 221.6 h, 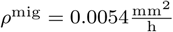, **2)** *τ*^rep^ = 206.7 h, *τ*^death^ = 90.5 h, *τ*^mut^ = 219.5 h, 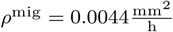, **3)***τ*^rep^ = 184.2 h, *τ*^death^ = 325.4 h, *τ*^mut^ = 186.5 h, 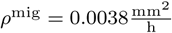, **4)** *τ*^rep^ = 233.5 h, *τ*^death^ = 143.0 h, *τ*^mut^ = 175.9 h, 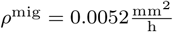, **5)** *τ*^rep^ = 201.2 h, *τ*^death^ = 295.5 h, *τ*^mut^ = 132.3 h, 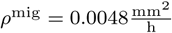, **6)** *τ*^rep^ = 219.8 h, *τ*^death^ = 177.2 h, *τ*^mut^ = 87.33 h, 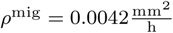. Other parameters are listed in Table 1.

### Evolutionary dynamics showed development of heterogeneity and dominance of the most aggressive clones

The simulator can be used to study the evolutionary dynamics within the tumor. An example is shown in Fig 5A, which shows the three-dimensional reconstruction of the tumor at three time points. In the first, the tumor mainly comprises the wild-type subpopulation, with small contributions of early-arising subtypes containing only one mutation. T the second time point we observe the emergence of more complex genotypes, with a significant contribution from the RTK mutated type, due to the effect of this mutation on proliferation. The end time point shows increased heterogeneity of the tumor with more altered genotypes taking over most of the tumor surface. This leads to the appearance of explosive peripheral areas (also resembling the lower parts of the tumor shown in Fig 4A). These features can be more clearly perceived in Fig 5B and Fig 5C, a reconstruction of the whole tumor in its final stages. In this case tumor diameter is around 7 cm, again in the range of real GBMs [28, 34]. It is important to point out that the three-dimensional reconstructions are isosurfaces of the total volume occupied by a subpopulation. There is overlapping, since two subpopulations or more can coexist in a given region of the tumor, hence the superposition of colors seen in the reconstruction. Fig 5D shows a phylogenetic reconstruction of the tumor. This is typically done in genetic analysis of GBM [19, 20]. Here we show that it is possible in the model to track the time at which a given mutation appeared and reconstruct from which population it came. Note that phylogenetic tree reconstructions of clonal lineages for individual GBMs have been performed by combining bulk exome sequencing with single-cell RNA-seq data [50].

**Fig 5.**
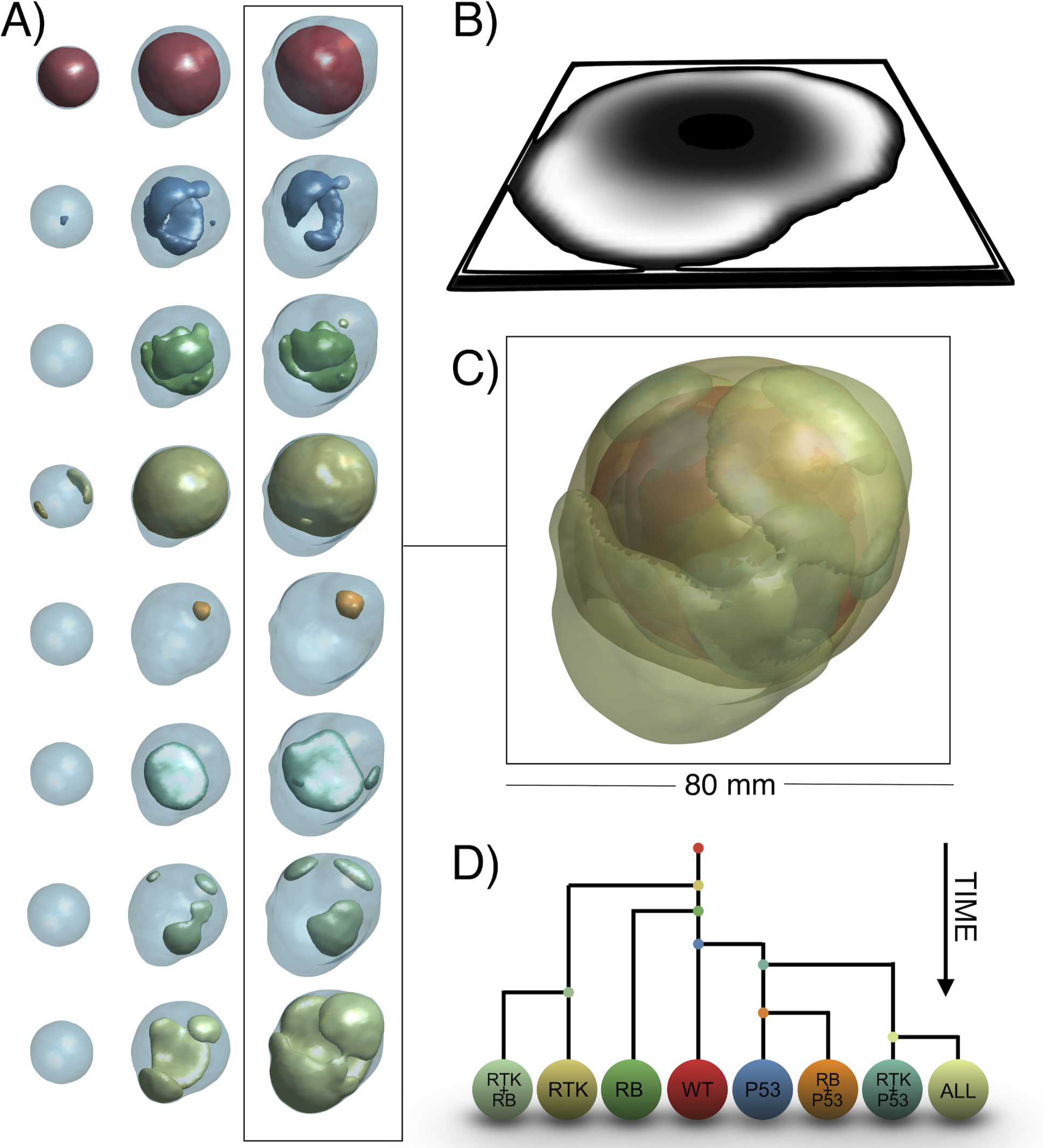
Example of the dynamics of the tumor’s clonal composition. **A)** Evolution of the eight clonal populations included in the model (one per row). Total tumor volume is shown as a light blue background. Times correspond to 8.5, 10 and 11.5 months. Parameters for this simulation are *τ*^rep^ = 179.1 h, *τ*^death^ = 292.5 h, *τ*^mut^ = 222.7 h, 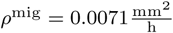. Other parameters are as in Table 1. **B)** Tumor central slice showing in white high tumor cell density. **C)** Three-dimensional subtype composition of the tumor. **D)** Reconstruction of the phylogeny of the tumor. Each bifurcation represents a mutation. Bifurcations occurring first in each branch represent mutations appearing earlier in time.

As new clonal subpopulations appear in the tumor, heterogeneity is expected to increase. However, due to cell competition and selection of the fittest, more aggressive subpopulations may end up occupying all the available space, confining less aggressive subpopulations to the core of the tumor and preventing them from proliferating, thus driving them to extinction. This fixation effect may result in a decrease in heterogeneity, as the fittest subpopulation becomes dominant. Fig 6A shows an example of the oscillatory behavior of the heterogeneity, as measured by the Shannon and Simpson’s indexes. Changes in these indexes are associated with proliferation of several subtypes or the dominance of one, respectively. This is more clearly seen in Fig 6B, which depicts the abundance of each subtype on a logarithmic scale. In this example simulation there is a clear increase in heterogeneity between months seven and ten of the simulation, with the coexistence of the first four subtypes. The dominance of the RTK subtype then causes heterogeneity to decrease, only to rise again with the appearance of more complex genotypes. Expansions and extinctions seen in this figure are compatible with the reconstructions shown in Fig 5A. Depending on when and where new clonal subpopulations appear, each simulation will bring a different evolutionary dynamic. Often, the most aggressive population carrying all alterations prevails, but a coexistence of two or more subpopulations may also take place. A combined view of the final state for all simulations is shown in Fig 6C. The heatmap below (Fig 6D) displays a clearer view of the possible endpoints of a simulation.

**Fig 6.**
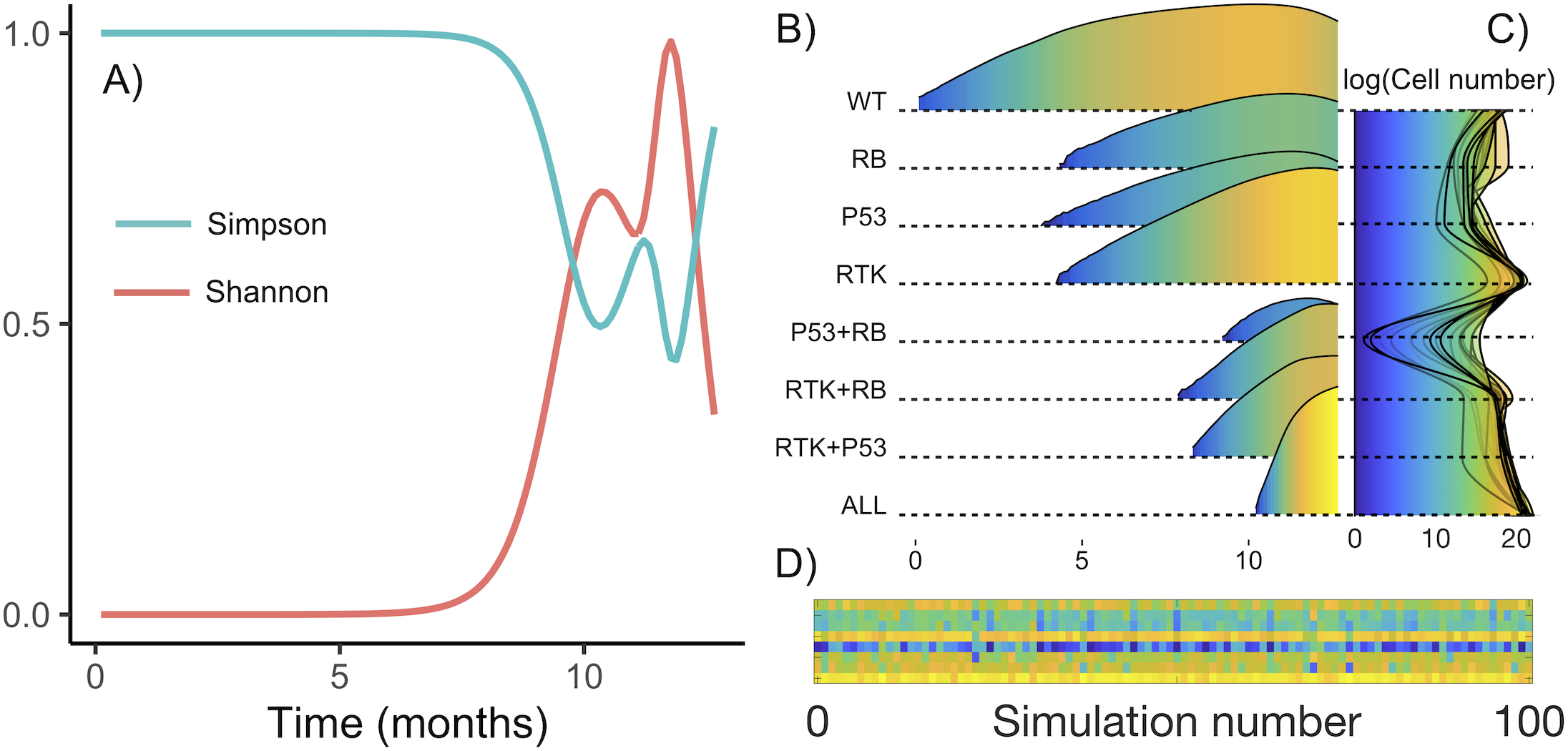
Dynamical behavior of tumor heterogeneity. **A)** Evolution of Shannon and Simpson’s indexes for a typical run. Basal rates for this run are *τ*^rep^ = 242.6 h, *τ*^death^ = 213.3 h, *τ*^mut^ = 221.6 h, 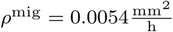. The other parameters are those of Table 1. **B)** Abundance of each subtype in logarithmic scale as a function of time. **C)** Superposition of final subtype abundance for all simulated tumors. **D)** Final abundance per subtype per simulation. Each row corresponds to one subtype, in the same order as above. Color indicates end-point abundance.

### Surrogates of tumor growth obtained from the model replicate the behavior of real GBMs

Much effort has been directed towards finding imaging-based prognostic biomarkers in GBM [16, 22, 23, 34, 51–55]. Our simulations allow the whole tumor natural history to be reconstructed, from its inception to the patient’s death. We focus our attention here on variables that can be obtained from our simulations and that have been found to have a prognostic value: Rim width, surface regularity and scaling law exponent (See ‘Methods’).

One of the most typical analyses in terms of prognostic value involves extracting the values of these parameters from diagnostic images and correlating them with patient survival. In our case, the model allows for tracking the evolution of these metrics over the whole tumor lifespan. Results for rim width and surface regularity for 100 simulations are shown in Fig 7. Time units have been normalized in order to compare simulations with different ranges of time evolution (see Fig 3). Curves represent the progression of the tumor from 1 cm^3^ to 100 cm^3^. Rim width typically increased with time as the tumor became more aggressive. This is an important difference with models having a simple clonal composition [49, 56], where the rim width was found to be constant. However, both approaches led to the same conclusion, namely, that rim width on diagnosis was associated with prognosis. Surface regularity was found to correlate with tumor clonal composition. Tumor slices corresponding to high and low values of each measure are also shown in Fig 7 to provide insight into their meaning.

**Fig 7.**
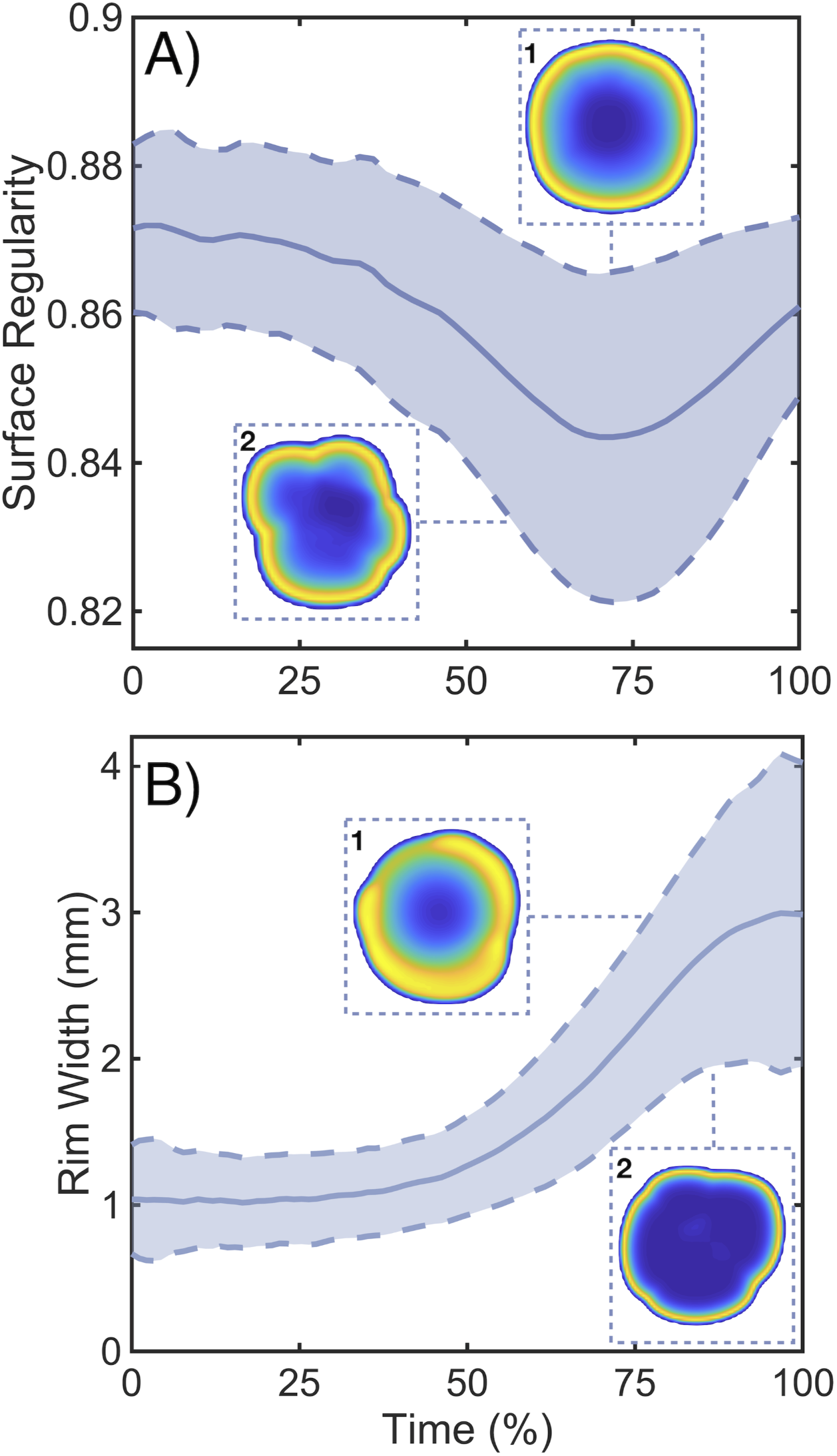
Dynamics of prognostic measures obtained from the model. **A,B)** Time evolution of spherical rim width and surface regularity for 100 simulations with parameters from Table 1. The solid line is the average value and the dashed line the standard deviation. 2D reconstructions correspond to characteristic upper and lower values of each variable. Basal parameters, measured in hours, for each subplot are: **A1)** *τ*^rep^ = 94.2 h, *τ*^death^ = 170.8 h, *τ*^mut^ = 99.4 h, 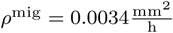, **A2)** *τ*^rep^ = 184.9 h, *τ*^death^ = 230.1 h, *τ*^mut^ = 120.0 h, 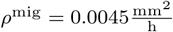, **B1)** *τ*^rep^ = 104.8 h, *τ*^death^ = 323.5 h, *τ*^mut^ = 222.7 h, 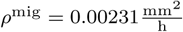, **B2)** *τ*^rep^ = 158.7 h, *τ*^death^ = 54.2 h, *τ*^mut^ = 197.7 h, 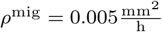.

As a test of the ability of our modeling methodology to replicate real tumor behavior we studied the association between the measures obtained from the simulations and overall survival, *in silico*. Since our model did not include therapy, typical survival times are expected to be short. A summary of survival analysis results is shown in Fig 8, where it is clear that all measures showed statistically significant curve separation. Median differences were found to be small, due to lack of treatment, but highly significant. These results indicate that surface regularity, rim width and scaling law exponent had prognostic value *in silico*, as happens in real tumors [21–23, 43]. Poor prognosis was associated with larger rim widths, lower surface regularity and larger scaling exponents.

**Fig 8.**
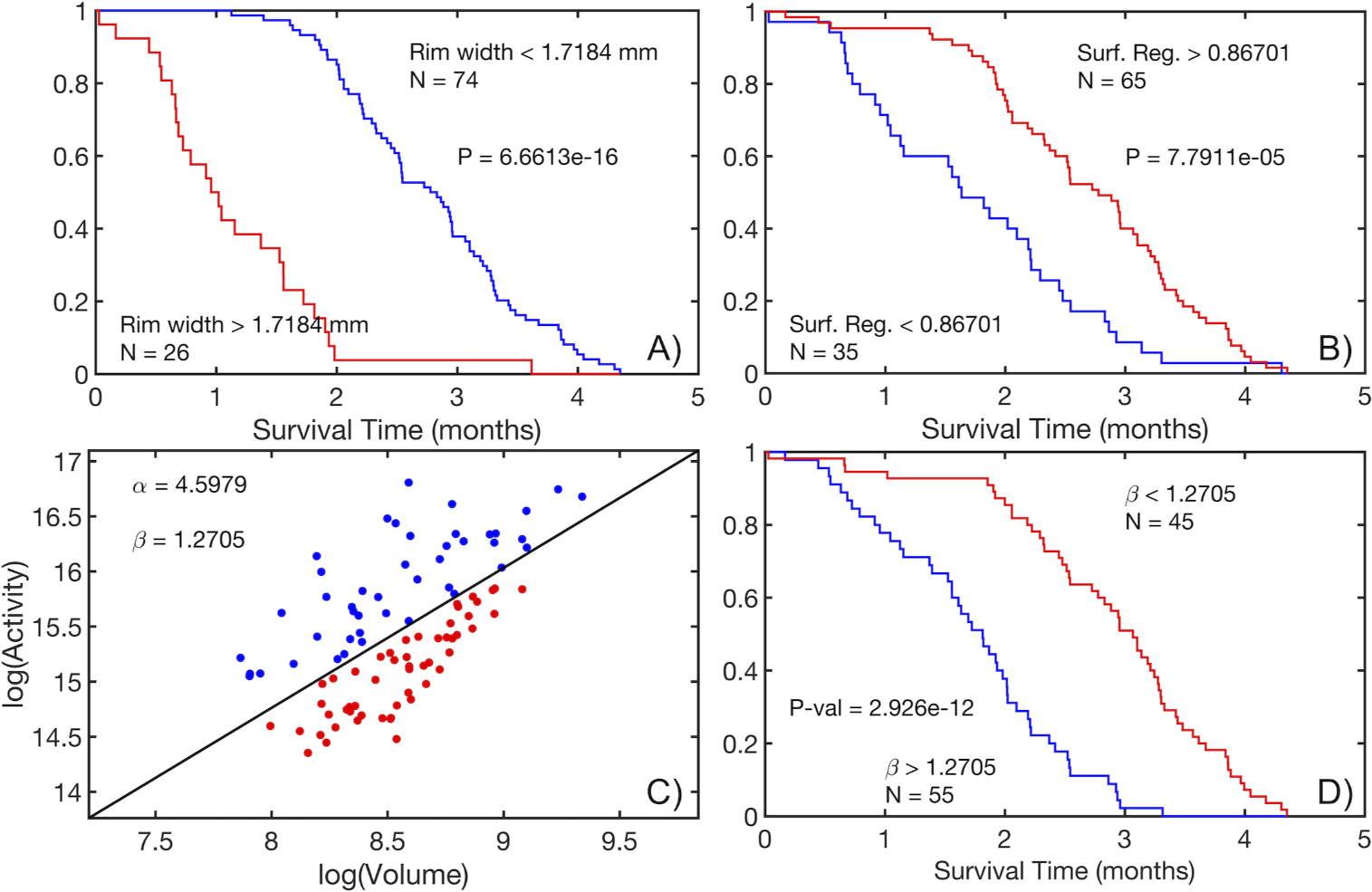
Prognostic measures obtained from the mathematical model recapitulate the behavior of those obtained from MRI and PET images of GBMs. **A)**. Kaplan-Meier curves for the population splitting using the spherical rim width taking a threshold equal to 1.7184 mm. Median survival difference between groups was 1.67 months. **B)**. Kaplan-Meier curves for the population splitting using the surface regularity taking a threshold equal to 0.86701. Median survival difference between groups was 0.67 months. **C)**. Scaling law exponent computation representing hypoactive (red) and hyperactive (blue) tumors. Parameters from linear regression from Eq (15) are shown. **D)**. Kaplan-Meier curves for the population splitting using the scaling law exponent, taking a threshold *β* = 1.2705. Median survival difference between groups was found to be 1.23 months.

### Mesoscopic simulation algorithm has good parallelization properties

One of the strong points of the mesoscopic simulation approach presented here is its low computational cost. Typical simulation times using the Julia code were a few minutes (3-5) in personal workstations and 80^3^ grid points. Although this is a remarkably small running time, there are several actions, such as increasing the lattice size, adding biological processes or performing many runs to explore parameter regions, that may lead to substantially longer computation times by orders of magnitude. This is why it is interesting to analyze the parallelization properties of the simulation algorithm under study.

Since grid points are updated synchronously, our algorithm has the potential for good scalability and thus benefit substantially from parallel computing. We performed a simple set of tests with our Julia implementation. Major bottlenecks were I/O to files and the main spatial loop, which iterates along all occupied voxels in order to update their cell numbers. While the former hardly benefits from parallelization, dividing the latter into several threads would improve run time. However, data communication between threads introduced a new bottleneck, which could reduce parallelization gains as the number of cores used increases.

We built a simple parallel version of the main code, and ran it with 1, 2, and 4 parallel threads, to check potential run time gains. Simulation runs with 2 cores took nearly 0.5 times less time to finish, while simulation runs with 4 cores improved run time by 60 %. This shows that our algorithm has the potential to benefit from parallel implementation in scenarios were it might be necessary do so (Fig 9).

**Fig 9.**
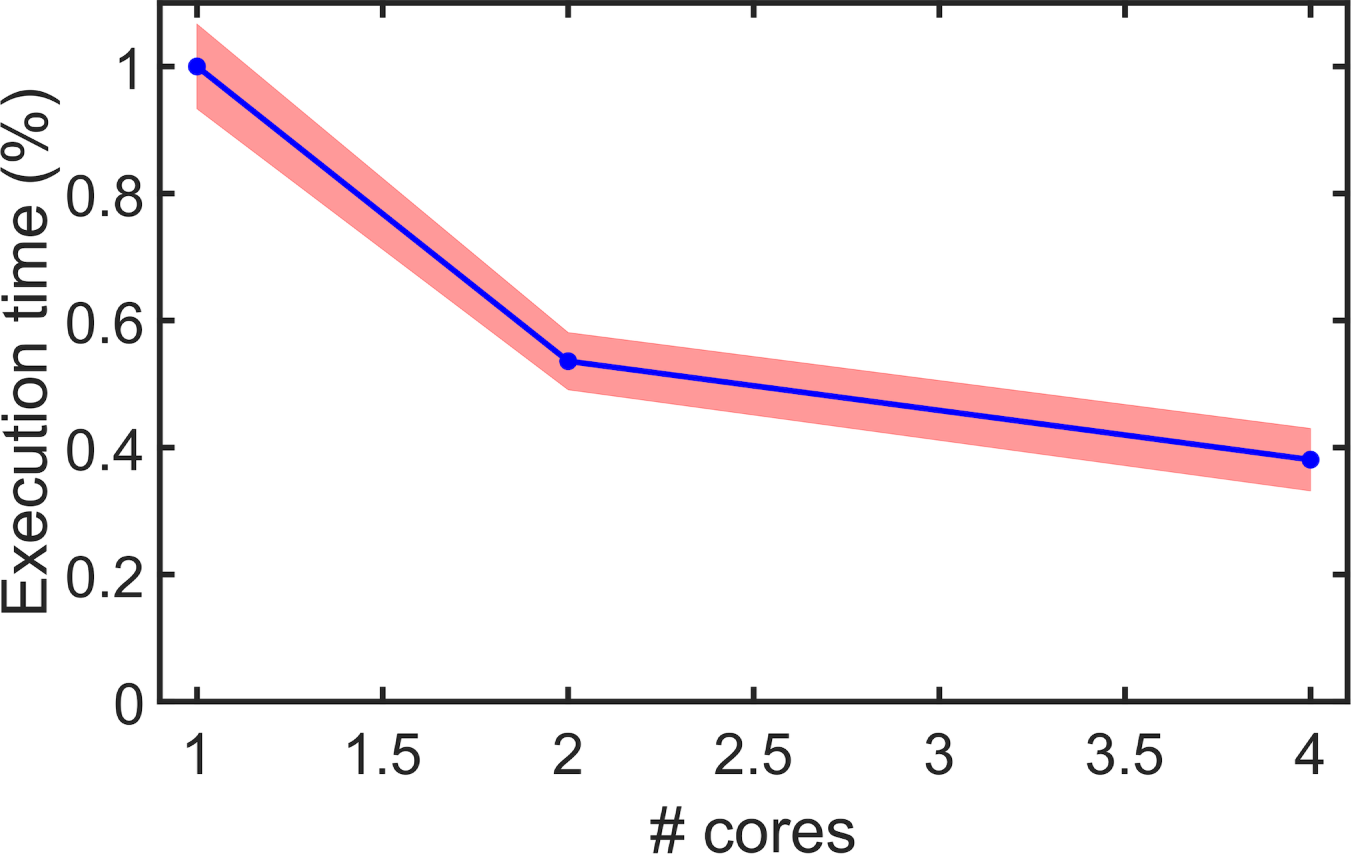
Code parallelization. Execution time of test cases as a function of the number of cores. Time is expressed as a percentage of single-core parallel version run time. Red band represents standard deviation.

## Discussion

In this paper we have presented a stochastic mesoscale simulator aimed at mimicking the natural history of a tumor from its inception to clinically observable sizes. Many different discrete simulation approaches are available to accomplish that task and shed light on different processes in oncology [1–6]. In our case the focus was on finding a balance between computational complexity and biologically meaningful assumptions, allowing for the study of a number of macroscopic features over the whole lifespan of the malignancy.

The tumor was described at the mesoscopic scale as a composition of different clonal populations at the voxel level, each of them having clone-specific characteristics that determine their behavior. Cells grew by proliferation, migrated and diversified as a result of mutational events that altered specific cellular processes. Death accounted for cell turnover and necrotic core formation. These biological rules were implemented as probabilistic events, incorporating both internal and external influences.

From the point of view of simulation, these rules were set up on a discrete three-dimensional space, following the perspective of multi-compartmental cell automaton models and matching the spatial resolution to current high-resolution medical imaging standards. The three-dimensional lattice was updated synchronously, taking the initial cell to a fully grown, spatially heterogeneous tumor. With efficiency in mind, the goal of this setup was to minimize computational time. This makes it possible to use our modeling framework to rapidly study different tumor dynamics scenarios and to test novel hypotheses. Other discrete modeling paradigms like those based on the Gillespie algorithm [57] provide correct solution trajectories for stochastic processes but become inefficient and computationally intensive when the number of events is high, due to the tau-leaping updating method. Finally, the algorithm allows for further improvements in speed by adding parallelization. Special emphasis was placed on the generation of a context where competition led to natural selection, a contribution framed in the mathematical modeling of evolutionary processes [58–60], rather than accurately parameterizing the model.

Dynamical behavior of the tumor was first analyzed in terms of volume and radial growth. Despite the logistic nature of each voxel’s dynamics, the whole tumor showed sustained growth, first linear and then accelerating as a result of the diversification and interplay of the populations, in a process that selects for more aggressive clones. Curve fitting resulted in power law being the most accurate description, over other unbounded laws like exponential or linear radial, pointing towards a relationship between metabolic activity, evolutionary dynamics and aggressivity [43]. The dynamics of simulated tumors changed from run to run as a result of the stochastic nature of the model, which allowed the influence of one-off events and parameter variability to br studied. This also benefited from the discrete consideration of the different variables, in line with previous suggestions [61]. The model provided a framework to analyze situations of clonal evolution of populations that are otherwise inferred from measurements of mutational spectrum and proportions [18–20, 62]. Furthermore, the capacity to extend this longitudinal simulation up to clinical sizes permits comparison of the dynamics with medical imaging, which can in turn be used in model calibration and quantitative description [63]. This opens the door to the analysis of other features such as the expansion and size of the necrotic core and its dependence on cell death and turnover.

Population diversification is another fundamental feature of the simulation and is analyzed here from an evolutionary ecology point of view, with metrics like the Shannon and Simpson indices [33]. This has already been done in heterogeneity analysis of tumors [64, 65]. The observed oscillations in such indexes reveal the process of clonal evolution and population fixation, in which aggressive phenotypes progressively displace the previous clones. Heterogeneity decreases when this happens and is maintained when competition is active in different regions of the tumor. This is interesting per se since heterogeneity is also spatially distributed, meaning that there may be spatial areas of the tumor with more diversity than others. A whole-tumor measure of diversity misses these characteristics. Three-dimensional reconstructions and exploration of the distribution of clonal populations enables the exploration of such scenarios. Again, the possibility of understanding this process longitudinally can be a source of hypothesis testing, particularly when combined with RNA-sequencing techniques to reconstruct phylogenetic trees of clonal lineages for individual GBMs. Moreover, our framework can readily incorporate the action of chemotherapeutic agents and capture the emergence of the different processes contributing to drug resistance [66].

Using the model, we studied three macroscopic quantities that have been proven to show prognostic value: surface regularity, tumor rim width and scaling exponent [21–23, 43]. The model simulations were able to reproduce the behavior of these significant metrics having clinical value. Time evolution curves of the first two variables showed a progressively increasing rim width up to a saturation value and a decrease in sphericity. Both can be explained as a result of the appearance of more malignant clones that take over at the boundary, giving way to a larger infiltrative area which manifests geometrically as a lobule protruding from the main tumor mass. This association between degree of malignancy, lobular shape and infiltrative area was confirmed by the survival analysis of both variables. With respect to the scaling exponent, the fit shows alignment with a previous study on the impact and emergence of the associated scaling law [43] and Kaplan-Meier analysis with the splitting threshold given by the exponent also shows a significant association with prognosis.

As with any simplified dynamical model in the biological sciences, in its simplicity lie both its virtues and its drawbacks. The downsides are the artifacts that appear during early stages of growth as a result of the discretization and its impact on the migration process; the absence of a microenvironment, its constituents and their influence on cellular processes and the lack of a clear distinction of genotype and phenotype in order to study the connection between them. Also, the number of clones is specified beforehand, in contrast to other evolutionary approaches in which the main elements emerge from a more simplistic consideration of features (see [9] for a comprehensive review). These drawbacks give us the future lines of work with the mesoscale simulator. At the same time it is used in research into evolutionary processes, efforts are to be directed towards improving its capabilities, especially the addition of external components such as the distribution of nutrients, their diffusion and the development of a pathophysiological vasculature. This is of utmost importance since many therapies are critically dependent on the tumor vasculature status. On the computational side, alternative forms of discretization and iterative configuration could be attempted. A last line of work is to develop methods for adjusting model parameters to specific situations and for estimating the impact of mutations/phenotypical changes on the cellular processes considered.

We hope this new methodology will be found to be a useful addition to the plethora of discrete simulation approaches intended to benefit cancer patients through the tools that computational approaches can provide.

## References

1. Deisboeck TS, Wang Z, Macklin P, Cristini V. Multiscale cancer modeling. Annual Review of Biomedical Engineering. 2011;13:127–155. https://doi.org/10.1146/annurev-bioeng-071910-124729 PMID: 21529163.

2. Rejniak KA, Anderson AR. Hybrid models of tumor growth. Wiley Interdisciplinary Reviews: Systems Biology and Medicine. 2011;3(1):115–125. https://doi.org/10.1002/wsbm.102 PMID: 21064037.

3. Wang Z, Butner JD, Kerketta R, Cristini V, Deisboeck TS. Simulating cancer growth with multiscale agent-based modeling. In: Seminars in Cancer Biology. vol. 30. Elsevier; 2015. p. 70–78. https://doi.org/10.1016/j.semcancer.2014.04.001 PMID: 24793698.

4. Metzcar J, Wang Y, Heiland R, Macklin P. A review of cell-based computational modeling in cancer biology. JCO Clinical Cancer Informatics. 2019;3:1–13. https://doi.org/10.1200/CCI.18.00069 PMID: 30715927.

5. Letort G, Montagud A, Stoll G, Heiland R, Barillot E, Macklin P, et al. PhysiBoSS: a multi-scale agent-based modelling framework integrating physical dimension and cell signalling. Bioinformatics. 2019;35(7):1188–1196. https://doi.org/10.1093/bioinformatics/bty766 PMID: 30169736.

6. Bravo RR, Baratchart E, West J, Schenck RO, Miller AK, Gallaher J, et al. Hybrid Automata Library: A flexible platform for hybrid modeling with real-time visualization. PLoS Computational Biology. 2020;16(3):e1007635. https://doi.org/10.1371/journal.pcbi.1007635 PMID: 32155140.

7. Fadai NT, Baker RE, Simpson MJ. Accurate and efficient discretizations for stochastic models providing near agent-based spatial resolution at low computational cost. Journal of the Royal Society Interface. 2019;16(159):20190421. https://doi.org/10.1098/rsif.2019.0421 PMID: 31640499.

8. Waclaw B, Bozic I, Pittman ME, Hruban RH, Vogelstein B, Nowak MA. A spatial model predicts that dispersal and cell turnover limit intratumour heterogeneity. Nature. 2015;525(7568):261–264. https://doi.org/10.1038/nature14971 PMID: 26308893.

9. Chowdhury D, Stauffer D. Evolutionary ecology *in silico*: Does mathematical modelling help in understanding “generic” trends? Journal of Biosciences. 2005;30(2):277–287. https://doi.org/10.1007/bf02703709 PMID: 15886463.

10. Hanahan D, Weinberg RA. The hallmarks of cancer. Cell. 2000;100(1):57–70. https://doi.org/10.1016/s0092-8674(00)81683-9 PMID: 10647931.

11. Celiku O, Gilbert MR, Lavi O. Computational modeling demonstrates that glioblastoma cells can survive spatial environmental challenges through exploratory adaptation. Nature Communications. 2019;10(1):5704. https://doi.org/10.1038/s41467-019-13726-w PMID: 31836713.

12. Stupp R, Mason WP, Van Den Bent MJ, Weller M, Fisher B, Taphoorn MJ, et al. Radiotherapy plus concomitant and adjuvant temozolomide for glioblastoma. New England Journal of Medicine. 2005;352(10):987–996. https://doi.org/10.1056/NEJMoa043330 PMID: 15758009.

13. Cancer Genome Atlas Research Network. Comprehensive genomic characterization defines human glioblastoma genes and core pathways. Nature. 2008;455(7216):1061–1068. https://doi.org/10.1038/nature07385 PMID: 18772890.

14. Parsons DW, Jones S, Zhang X, Lin JCH, Leary RJ, Angenendt P, et al. An integrated genomic analysis of human glioblastoma multiforme. Science. 2008;321(5897):1807–1812. https://doi.org/10.1126/science.1164382 PMID: 18772396.

15. Brennan CW, Verhaak RG, McKenna A, Campos B, Noushmehr H, Salama SR, et al. The somatic genomic landscape of glioblastoma. Cell. 2013;155(2):462–477. https://doi.org/10.1016/j.cell.2013.09.034 PMID: 24120142.

16. Wang Q, Hu B, Hu X, Kim H, Squatrito M, Scarpace L, et al. Tumor evolution of glioma-intrinsic gene expression subtypes associates with immunological changes in the microenvironment. Cancer Cell. 2017;32(1):42–56. https://doi.org/10.1016/j.ccell.2017.06.003 PMID: 28697342.

17. Ciriello G, Cerami E, Sander C, Schultz N. Mutual exclusivity analysis identifies oncogenic network modules. Genome Research. 2012;22(2):398–406. https://doi.org/10.1101/gr.125567.111 PMID: 21908773.

18. Sottoriva A, Spiteri I, Piccirillo SG, Touloumis A, Collins VP, Marioni JC, et al. Intratumor heterogeneity in human glioblastoma reflects cancer evolutionary dynamics. Proceedings of the National Academy of Sciences. 2013;110(10):4009–4014. https://doi.org/10.1073/pnas.1219747110 PMID: 23412337.

19. Johnson BE, Mazor T, Hong C, Barnes M, Aihara K, McLean CY, et al. Mutational analysis reveals the origin and therapy-driven evolution of recurrent glioma. Science. 2014;343(6167):189–193. https://doi.org/10.1126/science.1239947 PMID: 24336570.

20. Kim H, Zheng S, Amini SS, Virk SM, Mikkelsen T, Brat DJ, et al. Whole-genome and multisector exome sequencing of primary and post-treatment glioblastoma reveals patterns of tumor evolution. Genome Research. 2015;25(3):316–327. https://doi.org/10.1101/gr.180612.114 PMID: 25650244.

21. Molina D, Pérez-Beteta J, Luque B, Arregui E, Calvo M, Borrás JM, et al. Tumour heterogeneity in glioblastoma assessed by MRI texture analysis: a potential marker of survival. The British Journal of Radiology. 2016;89(1064):20160242. https://doi.org/10.1259/bjr.20160242 PMID: 27319577.

22. Pérez-Beteta J, Martínez-González A, Molina D, Amo-Salas M, Luque B, Arregui E, et al. Glioblastoma: does the pre-treatment geometry matter? A postcontrast T1 MRI-based study. European Radiology. 2017;27(3):1096–1104. https://doi.org/10.1007/s00330-016-4453-9 PMID: 27329522.

23. Pérez-Beteta J, Molina-García D, Ortiz-Alhambra JA, Fernández-Romero A, Luque B, Arregui E, et al. Tumor surface regularity at MR imaging predicts survival and response to surgery in patients with glioblastoma. Radiology. 2018;288(1):218–225. https://doi.org/10.1148/radiol.2018171051 PMID: 29924716.

24. Martínez-González A, Calvo GF, Romasanta LAP, Pérez-García VM. Hypoxic cell waves around necrotic cores in glioblastoma: a biomathematical model and its therapeutic implications. Bulletin of Mathematical Biology. 2012;74(12):2875–2896. https://doi.org/10.1007/s11538-012-9786-1 PMID: 23151957.

25. Schönfisch B, de Roos A. Synchronous and asynchronous updating in cellular automata. BioSystems. 1999;51(3):123–143. https://doi.org/10.1016/s0303-2647(99)00025-8 PMID: 10530753.

26. Milo R, Jorgensen P, Moran U, Weber G, Springer M. BioNumbers – the database of key numbers in molecular and cell biology. Nucleic Acids Research. 2009;38(Database Issue):D750–D753. https://doi.org/10.1093/nar/gkp889 PMID: 19854939.

27. Stensjøen AL, Solheim O, Kvistad KA, Håberg AK, Salvesen ø, Berntsen EM. Growth dynamics of untreated glioblastomas in vivo. Neuro-oncology. 2015;17(10):1402–1411. https://doi.org/10.1093/neuonc/nov029 PMID: 25758748.

28. Wang CH, Rockhill JK, Mrugala M, Peacock DL, Lai A, Jusenius K, et al. Prognostic significance of growth kinetics in newly diagnosed glioblastomas revealed by combining serial imaging with a novel biomathematical model. Cancer Research. 2009;69(23):9133–9140. https://doi.org/10.1158/0008-5472.CAN-08-3863 PMID: 19934335.

29. Nachman MW, Crowell SL. Estimate of the mutation rate per nucleotide in humans. Genetics. 2000;156(1):297–304. PMID: 10978293.

30. International Human Genome Sequencing Consortium. Initial sequencing and analysis of the human genome. Nature. 2001;409(6822):860–921. https://doi.org/10.1038/35057062 PMID: 11237011.

31. Venter JC, Adams MD, Myers EW, Li PW, Mural RJ, Sutton GG, et al. The sequence of the human genome. Science. 2001;291(5507):1304–1351. https://doi.org/10.1126/science.1058040 PMID: 11181995.

32. De Vleeschouwer S. Glioblastoma. Codon Publications; 2017. http://dx.doi.org/10.15586/codon.glioblastoma.2017

33. Magurran AE. Measuring Biological Diversity. 1st ed. Oxford: Blackwell Publishing; 2004.

34. Pérez-Beteta J, Martínez-González A, Pérez-García V. A three-dimensional computational analysis of magnetic resonance images characterizes the biological aggressiveness in malignant brain tumours. Journal of the Royal Society Interface. 2018;15(149):20180503. https://doi.org/10.1098/rsif.2018.0503 PMID: 30958226.

35. Vicente AMG, Pérez-Beteta J, Amo-Salas M, Pardo FJP, Martín MV, Valencia HS, et al. 18F-Fluorocholine PET/CT in the Prediction of Molecular Subtypes and Prognosis for Gliomas. Clinical Nuclear Medicine. 2019;44(10):e548–e558. https://doi.org/10.1097/RLU.0000000000002715 PMID: 31306196.

36. Hatt M, Laurent B, Fayad H, Jaouen V, Visvikis D, Le Rest CC. Tumour functional sphericity from PET images: prognostic value in NSCLC and impact of delineation method. European Journal of Nuclear Medicine and Molecular Imaging. 2018;45(4):630–641. https://doi.org/10.1007/s00259-017-3865-3 PMID: 29177871.

37. Coroller TP, Agrawal V, Huynh E, Narayan V, Lee SW, Mak RH, et al. Radiomic-based pathological response prediction from primary tumors and lymph nodes in NSCLC. Journal of Thoracic Oncology. 2017;12(3):467–476. https://doi.org/10.1016/j.jtho.2016.11.2226 PMID: 27903462.

38. Hofheinz F, Lougovski A, Zöphel K, Hentschel M, Steffen IG, Apostolova I, et al. Increased evidence for the prognostic value of primary tumor asphericity in pretherapeutic FDG PET for risk stratification in patients with head and neck cancer. European Journal of Nuclear Medicine and Molecular Imaging. 2015;42(3):429–437. https://doi.org/10.1007/s00259-014-2953-x PMID: 25416633.

39. Aerts HJ, Velazquez ER, Leijenaar RT, Parmar C, Grossmann P, Carvalho S, et al. Decoding tumour phenotype by noninvasive imaging using a quantitative radiomics approach. Nature Communications. 2014;5(1):1–9. https://doi.org/10.1038/ncomms5006 PMID: 24892406.

40. Desbordes P, Ruan S, Modzelewski R, Pineau P, Vauclin S, Gouel P, et al. Predictive value of initial FDG-PET features for treatment response and survival in esophageal cancer patients treated with chemo-radiation therapy using a random forest classifier. PLoS One. 2017;12(3):e0173208. https://doi.org/10.1371/journal.pone.0173208 PMID: 28282392.

41. Tagliafico AS, Bignotti B, Rossi F, Matos J, Calabrese M, Valdora F, et al. Breast cancer Ki-67 expression prediction by digital breast tomosynthesis radiomics features. European radiology experimental. 2019;3(1):1–6. https://doi.org/10.1186/s41747-019-0117-2 PMID: 31414273.

42. Zhou Y, Ma XL, Pu LT, Zhou RF, Ou XJ, Tian R. Prediction of Overall Survival and Progression-Free Survival by the 18F-FDG PET/CT Radiomic Features in Patients with Primary Gastric Diffuse Large B-Cell Lymphoma. Contrast media & molecular imaging. 2019;2019. https://doi.org/10.1155/2019/5963607 PMID: 31777473.

43. Pérez-García V, Calvo GF, Bosque JJ, León-Triana O, Jiménez J, Pérez-Beteta J, et al. Universal scaling laws rule explosive growth in human cancers. Nature Physics. 2020 https://doi.org/10.1038/s41567-020-09XX-X

44. West GB, Brown JH, Enquist BJ. A general model for ontogenetic growth. Nature. 2001;413(6856):628–631. https://doi.org/10.1038/35098076 PMID: 11675785.

45. Guiot C, Degiorgis PG, Delsanto PP, Gabriele P, Deisboeck T S. Does tumor growth follow a “universal law”? Journal of Theoretical Biology. 2003;225(2):147–151. https://doi.org/10.1016/s0022-5193(03)00221-2 PMID: 14575649.

46. Gerlee P. The model muddle: in search of tumor growth laws. Cancer Research. 2013;73(8):2407–2411. https://doi.org/10.1158/0008-5472.can-12-4355 PMID: 23393201.

47. Benzekry S, Lamont C, Beheshti A, Tracz A, Ebos JM, Hlatky L, et al. Classical mathematical models for description and prediction of experimental tumor growth. PLoS Computational Biology. 2014;10(8):e1003800. https://doi.org/10.1371/journal.pcbi.1003800 PMID: 25167199.

48. Miller AM, Shah RH, Pentsova EI, Pourmaleki M, Briggs S, Distefano N, et al. Tracking tumour evolution in glioma through liquid biopsies of cerebrospinal fluid. Nature. 2019;565(7741):654–658. https://doi.org/10.1038/s41586-019-0882-3 PMID: 30675060.

49. Pérez-García VM, Calvo GF, Belmonte-Beitia J, Diego D, Pérez-Romasanta L. Bright solitary waves in malignant gliomas. Physical Review E. 2011;84(2):021921. https://doi.org/10.1103/PhysRevE.84.021921 PMID: 21929033.

50. Müller S, Liu SJ, Di Lullo E, Malatesta M, Pollen AA, Nowakowski TJ, et al. Single-cell sequencing maps gene expression to mutational phylogenies in PDGF-and EGF-driven gliomas Molecular Systems Biology. 2016;12(11):889. https://doi.org/10.15252/msb.20166969 PMID: 27888226.

51. Wangaryattawanich P, Hatami M, Wang J, et al. Multicenter imaging outcomes study of The Cancer Genome Atlas glioblastoma patient cohort: imaging predictors of overall and progression-free survival. Neurooncology 2015; 17(11):1525–1537. http://dx.doi.org/10.1093/neuonc/nov117 PMID: 26203066

52. Ingrisch M, Schneider MJ, Nörenberg D, et al. Radiomic Analysis Reveals Prognostic Information in T1-Weighted Baseline Magnetic Resonance Imaging in Patients With Glioblastoma. Invest Radiol. 52(6):360–366 (2017). http://dx.doi.org/10.1097/RLI.0000000000000349 PMID: 28079702

53. Cui Y, Ren S, Tha KK, Wu J, Shirato H, Li R. Volume of high-risk intratumoral subregions at multi-parametric MR imaging predicts overall survival and complements molecular analysis of glioblastoma. Eur Radiol 2017; 27(9):3583–3592. http://dx.doi.org/10.1007/s00330-017-4751-x PMID: 28168370

54. Lao J, Chen Y, Li ZC, Li Q, Zhang J, Liu J, Zhai G. A Deep Learning-Based Radiomics Model for Prediction of Survival in Glioblastoma Multiforme. Sci Rep. 2017; 7(1):10353. http://dx.doi.org/10.1038/s41598-017-10649-8 PMID: 28871110

55. Molina-García D, Vera-Ramírez L, Pérez-Beteta J, Arana E, Pérez-García VM. Prognostic models based on imaging findings in glioblastoma: Human versus Machine. Sci Rep 2019; 9:5982. https://doi.org/10.1038/s41598-019-42326-3 6461644

56. Pérez-Beteta J, Belmonte-Beitia J, Pérez-García VM. Tumor width on T1-weighted MRI images of glioblastoma as a prognostic biomarker: a mathematical model. Mathematical Modelling of Natural Phenomena. 2020;15:10. https://doi.org/10.1051/mmnp/2019022

57. Gillespie DT. Exact stochastic simulation of coupled chemical reactions. The Journal of Physical Chemistry. 1977;81(25):2340–2361. https://doi.org/10.1021/j100540a008

58. Nowak MA, Sigmund K. Evolutionary dynamics of biological games. Science. 2004;303(5659):793–799. https://doi.org/10.1126/science.1093411 PMID: 14764867.

59. Michor F, Iwasa Y, Nowak MA. Dynamics of cancer progression. Nature Reviews Cancer. 2004;4(3):197–205. https://doi.org/10.1038/nrc1295 PMID: 14993901.

60. Sottoriva A, Spiteri I, Shibata D, Curtis C, Tavaré S. Single-molecule genomic data delineate patient-specific tumor profiles and cancer stem cell organization. Cancer Research. 2013;73(1):41–49. https://doi.org/10.1158/0008-5472.CAN-12-2273 PMID: 23090114.

61. Durrett R, Levin S. The importance of being discrete (and spatial). Theoretical Population Biology. 1994;46(3):363–394. https://doi.org/10.1006/tpbi.1994.1032

62. Kim J, Lee IH, Cho HJ, Park CK, Jung YS, Kim Y, et al. Spatiotemporal evolution of the primary glioblastoma genome. Cancer Cell. 2015;28(3):318–l328. https://doi.org/10.1016/j.ccell.2015.07.013 PMID: 26373279.

63. Alfonso JCL, Talkenberger K, Seifert M, Klink B, Hawkins-Daarud A, Swanson KR, et al. The biology and mathematical modelling of glioma invasion: a review. Journal of the Royal Society Interface. 2017;14(136): 20170490. https://doi.org/10.1098/rsif.2017.0490 PMID: 29118112

64. Maley CC, Galipeau PC, Finley JC, Wongsurawat VJ, Li X, Sanchez CA, et al. Genetic clonal diversity predicts progression to esophageal adenocarcinoma. Nature Genetics. 2006;38(4):468–473. https://doi.org/10.1038/ng1768 PMID: 16565718.

65. Park SY, Gönen M, Kim HJ, Michor F, Polyak K. Cellular and genetic diversity in the progression of in situ human breast carcinomas to an invasive phenotype. The Journal of Clinical Investigation. 2010;120(2):636–644. https://doi.org/10.1172/JCI40724 PMID: 20101094.

66. á lvarez-Arenas A, Podolski-Renic A, Belmonte-Beitia J, Pesic M, Calvo GF. Interplay of Darwinian selection, Lamarckian induction and microvesicle transfer on drug resistance in cancer. Scientific Reports. 2019;9(1):9332. https://doi.org/10.1038/s41598-019-45863-z PMID: 31249353.

